# NeuVue: A scalable and customizable framework for electron microscopy proofreading

**DOI:** 10.1101/2022.07.18.500521

**Authors:** Daniel Xenes, Lindsey M. Kitchell, Patricia K. Rivlin, Hannah Martinez, Victoria Rose, Caitlyn Bishop, Rachel Brodsky, Brendan Celii, Justin Ellis-Joyce, Diego Luna, Raphael Norman-Tenazas, Devin Ramsden, Kevin Romero, Marisel Villafañe-Delgado, Forrest Collman, William Gray-Roncal, Jacob Reimer, Brock Wester

## Abstract

Connectomic reconstruction from large image volumes produces segmentation and synaptic-assignment errors that must be resolved to support downstream analyses. As datasets have grown larger and teams more distributed, proofreading has become a critical operational bottleneck. Workflows for proofreading and error correction have not scaled commensurately with connectomic data production and may not accommodate heterogeneous proofreader expertise and machine-generated candidate edits. New tools are therefore needed to organize, prioritize, and coordinate proofreading at volume scale. Here we present NeuVue, a task-management and prioritization framework that operationalizes proofreading through atomic, auditable tasks for individual and team review, multistage routing across proofreader cohorts, performance and volume-state tracking, and integration with community annotation, visualization, and analysis services. We report the use of NeuVue across two volumetric datasets, supporting scalable proofreading by over forty proofreaders and producing over fifty thousand edits. NeuVue provides a reproducible human-in-the-loop framework for generating, validating, and maintaining large connectomic datasets.

## Main

Large-scale electron microscopy (EM) imaging has enabled a shift from isolated small circuit reconstructions towards larger, dynamically curated connectomic volumes that span whole brains and cubic-millimeter mammalian volumes. Recent dataset releases include the adult Drosophila central brain hemibrain^1^, the whole-brain FlyWire connectome^2^, the multimodal MICrONS mouse visual cortex dataset^3^, and a petavoxel reconstruction of the human cortex^4^. This scaling challenge is now also a central goal of the NIH BRAIN Initiative Connectivity Across Scales (BRAIN CONNECTS) program, which aims to develop the research capacity and technical capabilities needed to generate wiring diagrams spanning entire brains across multiple scales, including projects pursuing whole-mouse-brain and other very large connectomic pipelines. Advances in imaging, alignment, and automated machine segmentation have enabled the generation of datasets at this scale^5, 6^; however, analysis-grade reconstructions still require substantial proofreading to resolve merge and split errors in the image segmentation, recover incomplete dendritic arbors, and ensure accurate synaptic assignments to neurons, all while maintaining consistent tracking of annotations and objects as data curation proceeds.^7-9^

This need has driven the development of collaborative environments for proofreading and annotation, including CATMAID^10^, VAST^11^, Knossos^12^, webKnossos^13^, NeuTu^14^, and FlyWire^15^, as well as more recent infrastructure developments such as the Connectome Annotation Versioning Engine (CAVE) for versioned proofreading and annotation-aware analysis.^16^ Together, these systems have shown that large-scale community proofreading is possible. However, as datasets become richer in accompanying annotations and automated services, the surrounding workflows have expanded beyond direct split-and-merge editing to include targeted validation, consensus review, annotation-guided triage, and interactions with external analysis tools.^17-19^ These trends raise logistical problems that are only partly addressed by existing systems: how to turn candidate errors into manageable review units, how to distribute them across proofreaders with different expertise, how to decide which candidates should be reviewed first, and how to coordinate follow-up review and adjudication on shared data. A complementary proofreading management layer is therefore needed within these larger proofreading campaigns to package operations into reproducible tasks and deploy them in an orchestrated manner across distributed users.

Here we present NeuVue, a workflow framework for connectomic proofreading that operationalizes large-scale review through coordinated task management, impact-aware prioritization, multistage routing across proofreader cohorts, task-level provenance and performance tracking, and integration with community tools and data platforms. Originally developed to support curation of the MICrONS “Minnie65” connectome^3^, NeuVue has evolved into a more general platform for distributed proofreading across community datasets. Existing proofreading systems support core annotation and correction capabilities, but differ in the extent to which they support task-oriented workflow orchestration across distributed users (Table 1). NeuVue addresses this gap by representing units of proofreading work as atomic, configurable tasks whose results can be ingested into a variety of proofreading backends, depending on the needs of the project. We describe how NeuVue supports scalable and distributed workflows for segmentation correction, annotation-guided review, and human-in-the-loop machine learning (ML) techniques.

**Table 1.** Core capabilities across several EM proofreading tools. Entries are deliberately conservative and reflect capabilities explicitly described in the cited papers or official documentation. Green checkmark indicates evidence of full support, yellow half-diamond indicates evidence of partial support, and red x-mark indicates no evidence of support for each of the listed capabilities.

| Tool | Web-based | Precomputed Support | Spatial Annotations | Split/Merge Editing | Collaborative/ Shared Session | User Groups/ Task Permissions | Priority Task Queue | Task Telemetry |
| --- | --- | --- | --- | --- | --- | --- | --- | --- |
| NeuVue | ✓ | ✓ | ✓ | ✓ | ✓ | ✓ | ✓ | ✓ |
| CATMAID | ✓ | ✗ | ◊ | ✗ | ✓ | ✓ | ✗ | ✗ |
| VAST | ✗ | ✗ | ✓ | ✓ | ✗ | ✗ | ✗ | ✗ |
| NeuTu | ✗ | ✗ | ✓ | ✓ | ✓ | ✗ | ✗ | ✗ |
| WEBKNOSSOS | ✓ | ✓ | ✓ | ✓ | ✓ | ✓ | ◊ | ✓ |
| FlyWire | ✓ | ✓ | ✓ | ✓ | ✓ | ◊ | ◊ | ◊ |

✓ Explicitly Supported
◊ Partially Supported
✗ Not Supported

## Results

### NeuVue packages proofreading as configurable, auditable tasks

NeuVue is organized around a task abstraction that couples a localized biological or segmentation question to the information needed to resolve it, including task instructions, priority, provenance, viewer state, and structured metadata [Fig. 1a]. This allows an atomic proofreading task type to be deployed across datasets, users, and review modes. For example, a task may ask whether a proposed split correctly separates two neuronal somata at a candidate merge site. In this case, NeuVue can present the proofreader with a **forced-choice task**, in which the user selects from a fixed set of predefined responses, such as accepting the proposed split, rejecting it, or marking the task as uncertain. This design enables rapid triage of large candidate sets before ambiguous cases are escalated to point-adjustment or direct-edit workflows. Tasks can be assigned directly to individuals, user groups, or anonymous public queues to enable flexible proofreading protocols. Proofreader groups can be configured independently and outputs from one queue can trigger follow-up tasks for adjudication, refinement, or consensus review.

**Figure 1.**
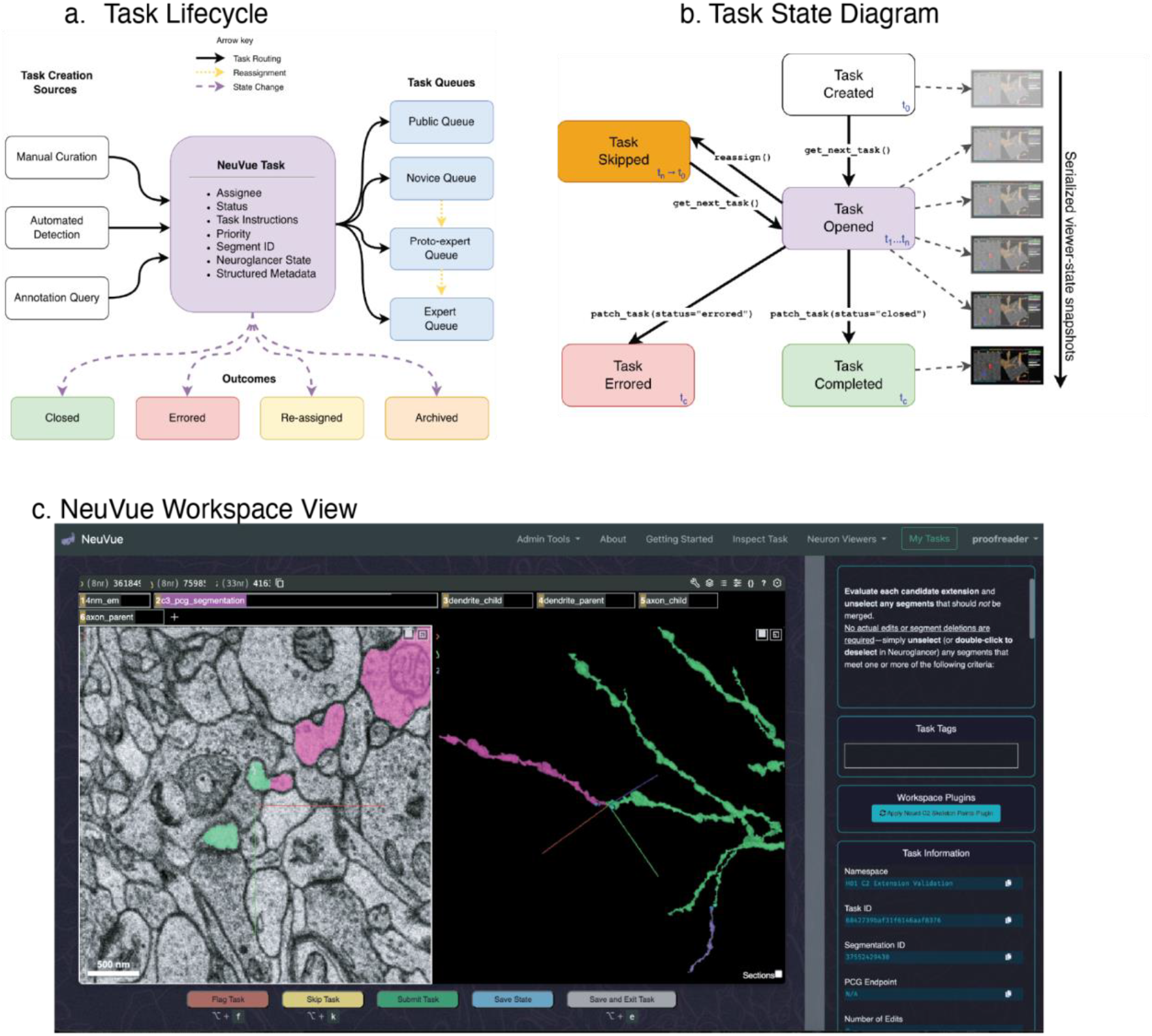
**(a)** A diagram of a task object, itemizing the minimum set of attributes needed to define a task. Tasks are sourced from manually curated errors, automated detection techniques such as NEURD, or through querying available annotation stores such as CAVE. Tasks can be assigned directly or to a group, exposed to a public queue, or chained through several proofreader groups for review workflows. **(b)** Task state transition diagram. NeuVue records a timestamped provenance entry whenever a task is created, opened, skipped, enters an error state, or is closed. On each task access and state transition, the current viewer state is saved, enabling a comprehensive and auditable changelog for the task. **(c)** NeuVue workspace user interface. Tasks are rendered in a configurable Neuroglancer distribution embedded within a webpage containing available proofreader actions and task information.

NeuVue records task state transitions, saved viewer states, and user actions as structured telemetry [Fig. 1b]. These logs provide provenance for individual decisions, support auditing and agreement analyses across proofreaders, and enable downstream automation such as queue prioritization, task chaining, and targeted retraining of error-detection services. In this way, NeuVue functions not only as a proofreading interface but as a workflow layer for reproducible dataset validation at community scale.

The NeuVue interface is configurable rather than tied to a single proofreading client [Fig. 1c]. A task can be rendered against custom visualization backends, including current and legacy Neuroglancer^20^ deployments. This broadens the range of workflows and data sources that NeuVue supports. To extend functionality without rebuilding the interface, we implemented a plugin system in which external analysis services can operate within the task context. For example, plugin services can call automated analysis pipelines such as the *NEURal Decomposition tool* (NEURD) to identify error regions, generate candidate corrections, or launch targeted follow-up tasks on demand.^9^ Integration with CAVE further enables tasks to query and display linked annotations, allowing users to inspect and act on nuclei, synapses, and cell-type predictions within the same proofreading environment.^16^

### NeuVue routes tasks according to proofreader expertise

NeuVue explicitly accommodates proofreader heterogeneity through group-based permissions, queue policies, and reassignment pathways. Rather than allowing unrestricted task selection, tasks are delivered according to administrator-defined priorities and administrators can organize users into groups that reflect proofreading proficiency and permitted actions. This allows low-risk validation tasks, direct edit tasks, and adjudication tasks to be routed to different users while preserving a common task abstraction and interface [Fig. 2a].

**Figure 2.**
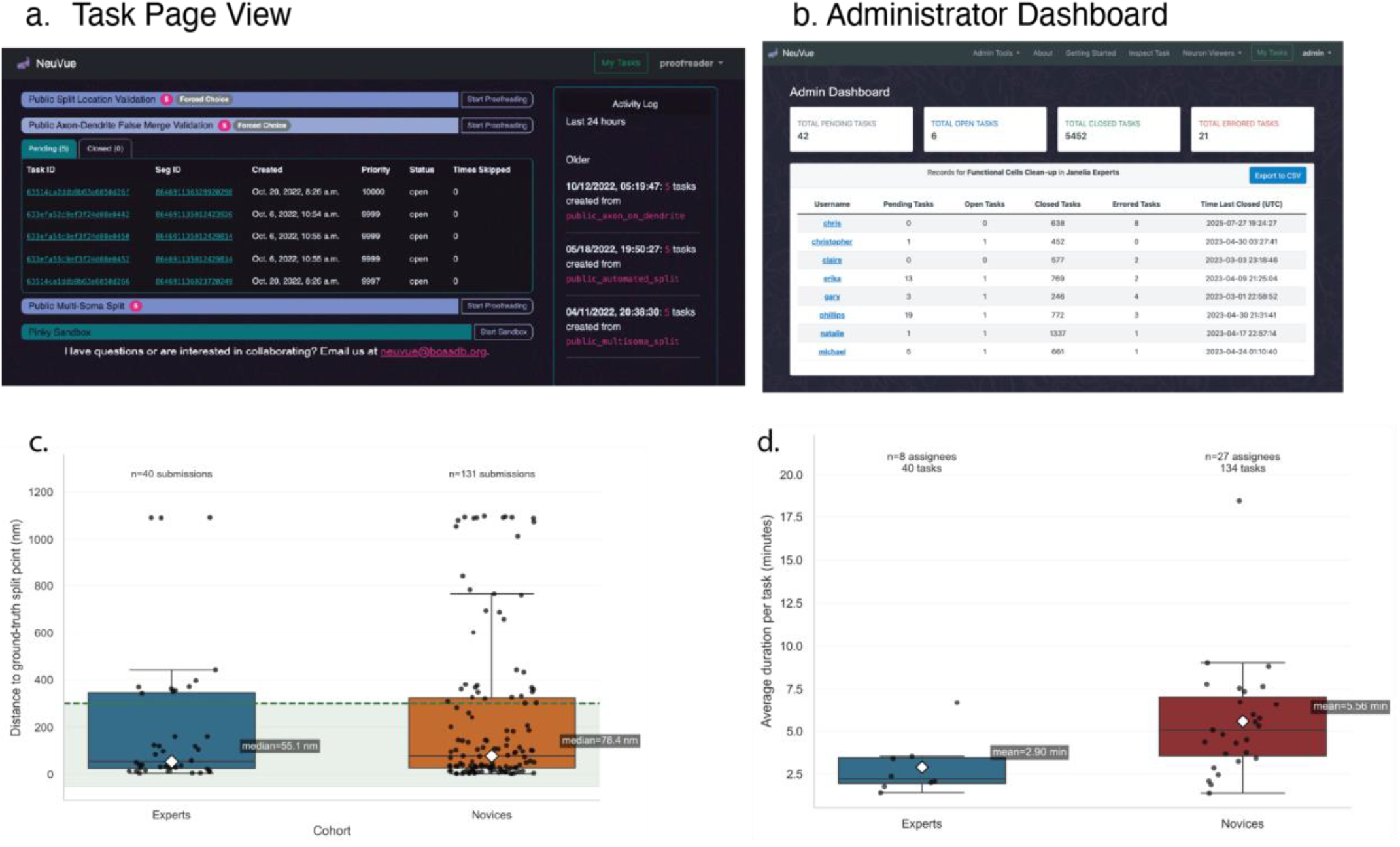
**(a)** Task page where proofreaders can select a task type to proofread. Tasks are sorted by activity and quantity. Tasks will also display the pending amount in the queue and a submission method label. **(b)** Administrator tools such as this dashboard view provide insight on completed tasks. Selecting a user allows the administrator to view that user’s task queue and inspect their results. **(c)** Distances from each proofreader’s submitted split point to the ground-truth split point for multi-soma benchmark tasks, shown by cohort (Experts, Novices). Points represent per-task distance across shared tasks. The dashed horizontal line marks the 300 nm agreement threshold; the shaded green region indicates within-threshold distances. While experts and novices both have high agreement rates with the ground truth tasks, experts are more consistently within threshold. **(d)** Average annotation time per task for Experts and Novices on multi-soma benchmark tasks. Each point is one proofreader’s mean task duration. Experts close tasks 47% faster than novices.

Administrators use dashboard and reporting tools to monitor progress and modify queue state at the level of individual users or groups [Fig. 2b]. In the Minnie65 proofreading campaign, this included reassigning, reprioritizing, and filtering tasks by task type, user, group, and time window, enabling active control over which proofreaders received which classes of work. Proofreaders can also self-manage parts of their workload by drawing from unassigned pools when appropriate for their experience level.

This routing framework enabled a staged review model. Novices contributed through constrained, low-risk decision tasks that also served as training, whereas ambiguous or edit-intensive cases were escalated to expert or proto-expert groups. In the semi-automated split workflow, forced-choice tasks allowed proofreaders to rapidly evaluate many candidate errors before making direct edits, and responses indicating that a proposed split was incomplete or misplaced automatically routed those cases to follow-up review or manual repair. Proofreaders with high agreement are promoted to a proto-expert group with direct write permissions and difficult multi-neuron IDs are pushed onward for expert review.

The resulting separation of roles is reflected in the task distribution and task submission style. Forced-choice review was performed at scale by both expert and novice cohorts, whereas edit-intensive queues concentrated in expert and proto-expert cohorts. Within the multi-neuron splitting workflow, lower-experience proofreaders were preferentially assigned easier two-soma cases (proofreading split), reserving more complex objects for expert review. This role-aware routing reduced unnecessary expert effort while maintaining a path for difficult decisions to receive additional scrutiny. We next asked whether this role-aware routing could be used to turn large numbers of automatically proposed split candidates into verified corrections and, ultimately, analysis-ready neurons.

To quantify differences between proofreader cohorts on the relevant proofreading goal of reducing multi-soma merges, we created a small multi-soma split benchmark dataset (n=5) with adjudicated ground-truth split points. We had novice and expert proofreaders complete the same tasks in their own respective queues. Proofreader agreement to the ground truth was defined by the Euclidean distance between a submitted split point and the ground truth split point, with placements within 300 nm counted as concordant. With this threshold, both cohorts, treated as a consensus average, submitted split points that agreed with ground truth on 4 of the 5 benchmark tasks; however, experts placed points closer to the adjudicated locations overall, with a median distance of 55 nm compared to 78 nm [Fig. 2c]. Experts also completed the benchmark more quickly: across assignee-level averages, the mean task duration was 2.90 min for experts and 5.56 min for novices [Fig. 2d]. This pattern is consistent with prior connectomic work showing that reconstruction errors differ in their downstream consequences and that proofreading effort is best directed toward the errors most likely to alter circuit interpretation^21^; in this setting, that rationale supports reserving ambiguous multi-soma merge corrections for expert review.

### Linked task groups convert candidate errors into analysis-ready neurons

We initially focused NeuVue on multi-soma errors because they were abundant, broadly distributed across the Minnie65 volume, and disproportionately disruptive to connectivity. In the starting materialization cell-type classification, 7,801 multi-soma segmentation IDs contained at least one neuronal soma, including 5,679 multi-neuron IDs and 2,122 neuron-plus-non-neuron merges.^22^ NEURD-generated candidate split sites then provided the first large semi-automated test of the platform: 10,148 candidate locations were supplied and 9,459 remained relevant after filtering stale sites from an older materialization.

To triage these candidates, we used a staged queuing strategy in which proofreaders first performed rapid forced-choice validation, with only ambiguous cases escalated to manual correction. This produced 22,522 expert forced-choice tasks, 5,139 novice forced-choice tasks, and 4,480 guided manual review tasks. Novice proofreaders contributed approximately 501 hours of first-pass review, allowing expert effort to be concentrated on ambiguous or edit-intensive cases.

After the semi-automated queues were depleted, NeuVue supported a shift to direct repair of the remaining high-impact merge errors. Approximately 3,500 multi-neuron IDs were placed into a task queue and more than 1,200 merged soma IDs were isolated into a separate manual workflow because they lacked reliable automated corrections. These queues yielded 1,740 expert review multi-neuron tasks, 1,606 proto-expert review multi-neuron tasks and 376 merged soma tasks. The same task abstraction therefore supported both rapid triage and longer expert interventions without refactoring the underlying proofreading environment.

At dataset-scale, these workflows reduced the number of multi-soma IDs containing at least one neuron by 6,614 and generated 15,632 new single-neuron IDs [Fig. 3]. Downstream analyses tracked 15,488 proofread neurons, and proofreading removed nearly all of the exceptionally large multi-neuron objects present in the initial materialization. The unresolved remainder was dominated by edge-adjacent objects, dense low-quality regions, and objects that were too large or unstable to edit reliably in the underlying CAVE datastack.

**Figure 3.**
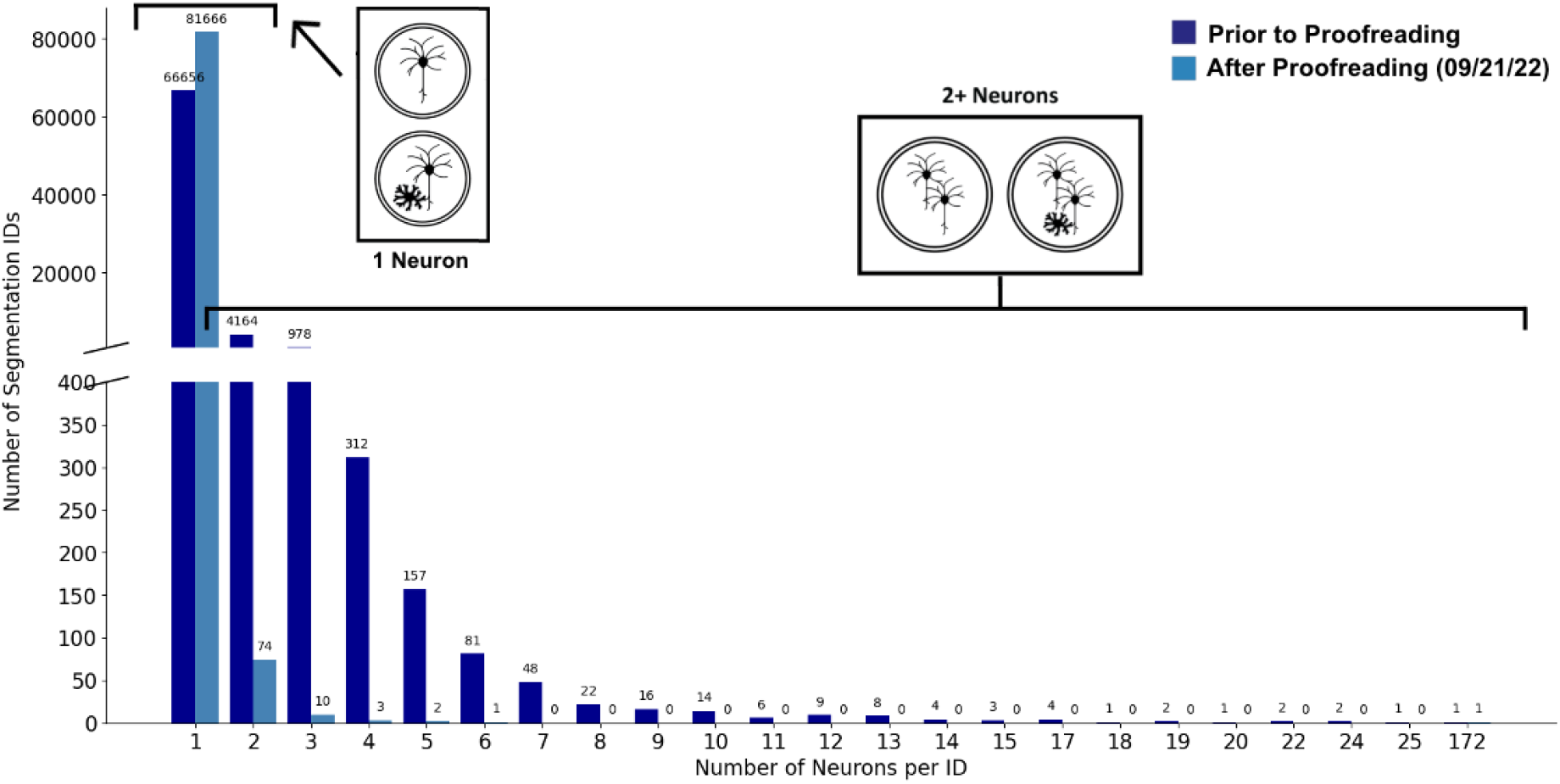
Difference in multi-neuron root IDs before (dark blue, materialization v272) and after NeuVue-enabled proofreading efforts up to 09/21/22 (lighter blue). Post proofreading illustrates a clear and significant reduction in multi-neuron root IDs in the volume.

We then aimed to transform newly split IDs into analysis-ready neurons rather than only correcting large segmentation errors. Of approximately 15,000 newly created single-soma IDs examined, about 1,500 fell outside predefined completeness thresholds and were placed into a targeted clean-up queue. This produced 584 single-soma clean-up tasks, totaling 119 hours of proofreading. After this second pass, 15,092 of 15,488 proofread neurons fell within the population-based completeness screening criteria, and 396 remained outside the acceptable range. Among 699 proofread neurons with functional co-registration, all satisfied the completeness criteria.

Across these workflows, NeuVue logged 36,447 completed tasks and 2,451 hours of proofreading time, corresponding to 6.31 generated single-soma neurons per hour overall and 6.15 single-soma neurons per hour for neurons that ultimately met our completeness thresholds. The platform also recorded 54,366 edits to the CAVE datastack, including 45,275 splits and 9,091 merges, linking queue design directly to edit burden and final reconstruction quality.

### NeuVue serves as the generalizable human-validation layer of CONNECTS-Proof

We next asked whether the NeuVue workflow framework would transfer beyond the Minnie65 campaign to a dataset with different automation and proofreading conditions. Within CONNECTS-Proof, an NIH BRAIN Initiative Connectivity Across Scales (BRAIN CONNECTS) project that combines NEURD-based automated error detection and correction with human validation and review for large-scale EM datasets, NeuVue serves as the interactive human-in-the-loop layer for candidate edits. The H01 dataset^4^ was chosen as the central dataset for this new approach, as it provided a high-impact, BRAIN CONNECTS–relevant mammalian test case that emphasized automated error detection and correction workflows run with a much smaller proofreader cohort.

We first asked whether NEURD-derived task metadata could be used to increase the value of limited expert review time. To do this, we implemented a queueing strategy in which each validation task was assigned a scalar priority based on its predicted connectomic impact, estimated from the number of synapse assignments expected to change if the proposed edit were accepted. In an experiment involving 104 forced-choice validation tasks, prioritized review increased the realized synaptic impact by 2.2-fold (95% CI, 1.42–3.58; n = 30 tasks) relative to random ordering. For the first 30 tasks, prioritized review affected 788 synapses, compared with a bootstrap mean of 246.0 synapses (95% CI, 171.9–325.2; n = 30 tasks) [Fig. 4a]. These results show that NeuVue can use NEURD outputs not only to generate tasks, but also to direct proofreaders toward the subset of machine-generated edits with the highest expected downstream value.

**Figure 4.**
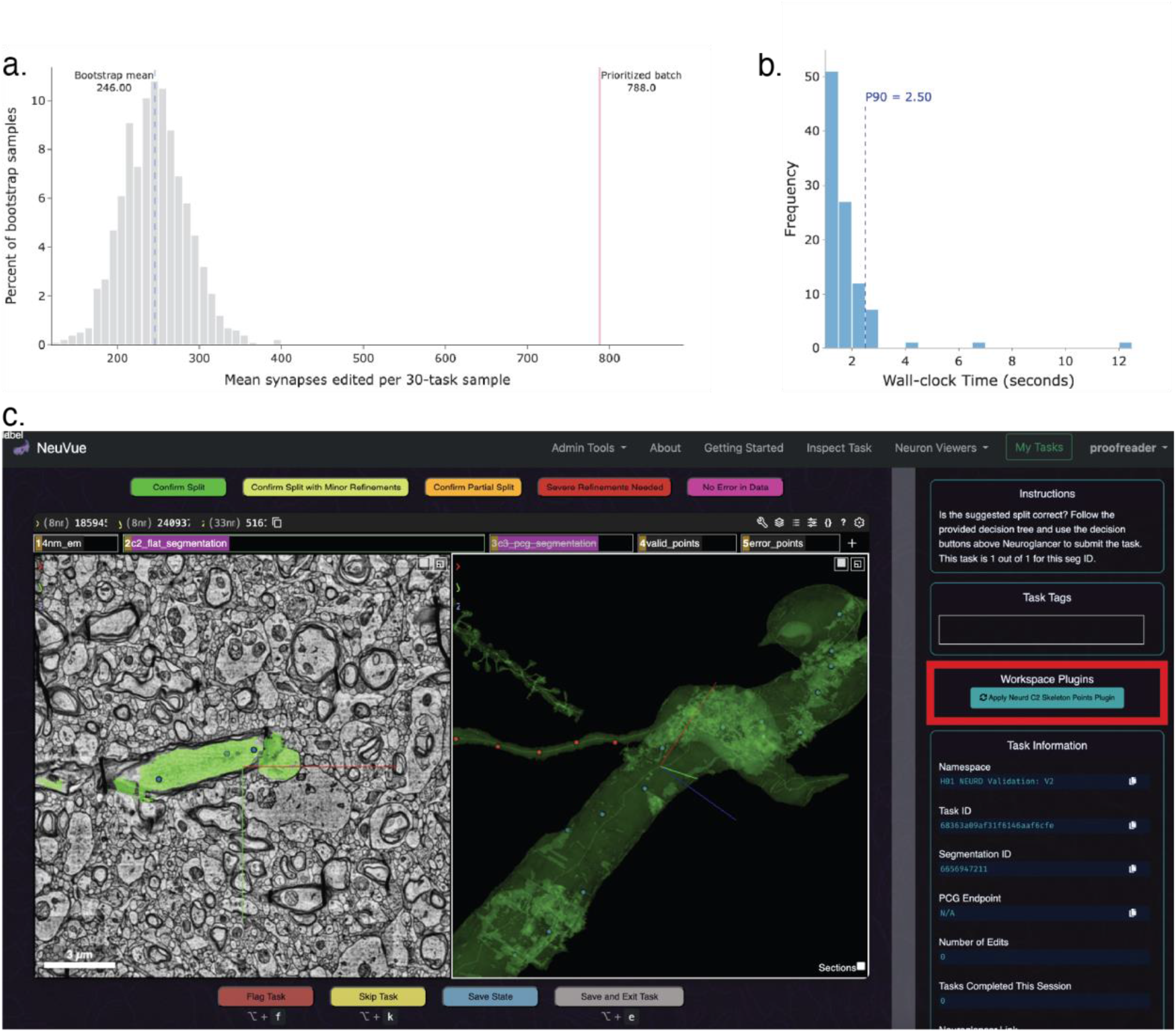
**(a)** Comparison of realized synapse-based impact for the prioritized first 30 split-validation tasks against the bootstrap distribution from randomly ordered samples of 30 tasks drawn from the same set of 104 proposed split edits. **(b)** Distribution of wall-clock times for on-demand pipeline invocation on 100 unique NeuVue tasks. 90th percentile or P90 latency measured at 2.5 seconds. **(c)** NeuVue proofreading workspace with an active plugin. When an on-demand plugin is available for a task type, the Workspace Plugins box on the right (highlighted in red) contains a button the user can click to trigger the service, allowing for up-to-date NEURD annotations to populate their Neuroglancer viewer.

We next extended NeuVue with an on-demand plugin system that allows the proofreading workspace to query NEURD services at task runtime. This design addressed a practical limitation of static task generation: when multiple edits affect the same cell, precomputed views can quickly become stale. In the proof-of-concept implementation, an on-demand plugin retrieved updated NEURD skeleton annotations for the cell currently under review and inserted them directly into the live workspace. Across 100 invocations, this service achieved a 90th percentile latency, or P90 latency, of 2.50 seconds [Fig. 4b]. This capability allowed proofreaders to evaluate edits against more current cell state without rerunning the full NEURD pipeline after every task. [Fig. 4c]

These improvements were directly used in the semi-automated validation of NEURD edits in the H01 dataset. A total of 449 validated merge (extension) edits and 939 validated split edits were reviewed in NeuVue before final filtering and bulk deployment. These task types supported the highest validation throughputs reported for this platform, with average task close rates of 0.66 ± 0.42 split-validation tasks per minute and 4.27 ± 3.16 extension-validation tasks per minute.

## Discussion

NeuVue suggests that workflow design is a distinct lever for scalable connectomics alongside improved segmentation models and richer proofreading interfaces. Originally developed within the IARPA MICrONS program, NeuVue demonstrates that large proofreading campaigns can combine structured training, cohort-based permissions, and staged escalation to coordinate heterogeneous proofreader teams at dataset scale. Across the Minnie65 and H01 proofreading campaigns, this framework supported 37,835 completed tasks, 2,493 hours of proofreading, and 54,366 recorded human edits. In Minnie65, expertise-aware routing concentrated expert effort on the most ambiguous and edit-intensive cases while allowing novice cohorts to absorb large first-pass validation queues; consistent with this design, experts completed matched multi-soma benchmark tasks 47% faster than novices and placed split points closer to adjudicated ground truth. In H01, priority ordering based on predicted connectomic impact increased the realized synaptic impact of early review by 220% relative to random ordering. These results argue that NeuVue’s main contribution is not a single proofreading workflow or algorithm, but an operational framework for turning distributed human review into reproducible, dataset-scale connectome curation.

Our results indicate that scalable proofreading depends as much on queue/task design as on the interface or data visualization layer alone. NeuVue tasks can be configured not only to ask whether an edit is correct, but also whether a candidate ranking is useful, whether a proposed edit is operationally deployable to the underlying CAVE datastack, and where automated error detection fails to surface the true correction at all. This flexibility matters because edit-by-edit validation does not fully capture practical performance of automated proofreading methods such as NEURD.

Several limitations exist within NeuVue and our broader proofreading approach. First, the Minnie65 results are dominated by high-impact merge correction and subsequent clean-up, so performance across newer task families remains less mature than for multi-soma splitting. Second, the main completeness assessment is population-based and anchored to previously proofread reference neurons; although this was practical at scale, it does not by itself comprehensively capture biological correctness or synapse-level error. Third, the remaining unresolved objects in Minnie65 were concentrated near dataset edges, in dense low-quality regions, or in exceptionally large objects that were difficult or impossible to edit in the underlying CAVE datastack, indicating that improved workflow design cannot fully compensate for upstream data limitations. Finally, several newer capabilities, including on-demand plugins and adaptive queueing, are currently supported by strong proof-of-concept results, but still need broader benchmarking across datasets, proofreader cohorts, and downstream analyses.

A natural next step is to evaluate these proofreading workflows with more explicit consensus and downstream quality metrics, including before-and-after synapse analyses and segmentation benchmarks such as variation of information and expected run length, while continuing to generalize task schemas to richer annotations and multimodal validation. If these directions hold, NeuVue could help shift proofreading from an ad hoc manual bottleneck toward a reproducible, modular component of large-scale connectomics production pipelines.

## Online Methods

### NeuVue Architecture

NeuVue comprises three interacting components: a cloud-hosted task database (NeuVue-Queue), a browser-based proofreading application, and administrator-facing tools for task creation, monitoring, and analysis [Fig. 5]. Administrators generate tasks through an API client and assign them to task namespaces corresponding to specific task types. Each task stores instructions, priority, author and assignee information, status, tags, visualization state, structured metadata, and timing telemetry. Tasks were designed to be atomic and stateless so that they could be completed independently and reordered at scale.

**Figure 5.**
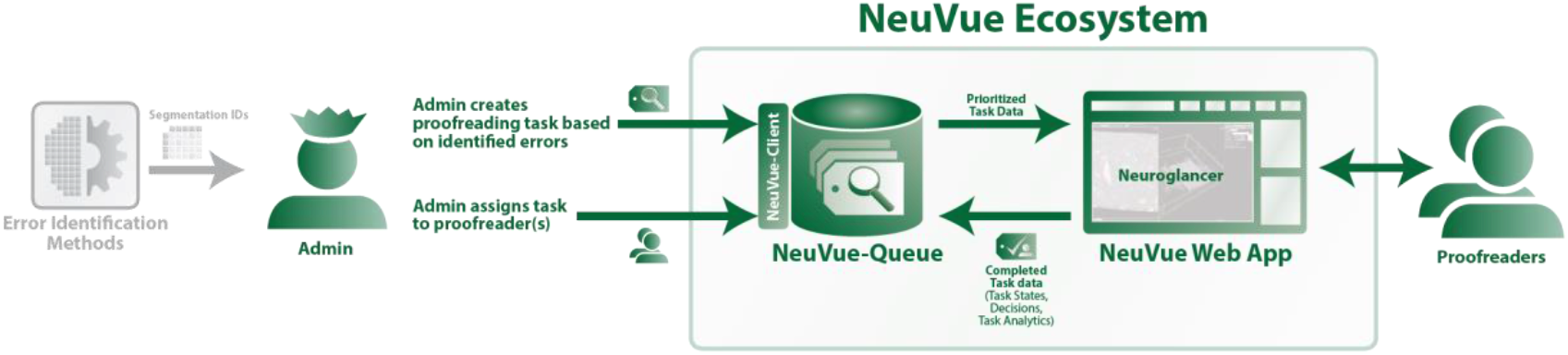
NeuVue component diagram. Segmentation IDs sourced from error detection methods are provided to NeuVue administrators to create individual tasks. NeuVue administrators create the task type based on the identified error information and assign it to proofreaders through the NeuVue-Queue API (implemented in NeuVue-Client). Proofreaders interact with the NeuVue Web App to view the Neuroglancer state and instructions associated with the task. The completed task data is saved back into the queue for later retrieval.

Users access NeuVue through authenticated browser sessions and work from task pages that summarize open, pending, and completed assignments. When a task is launched, the workspace loads a stored visualization state and task metadata into an embedded proofreading client. Proofreaders can complete tasks by committing direct split or merge edits, submitting forced-choice decisions or entering structured annotations, depending on the task definition. Queue actions, including skip, flag, save state, and save-and-exit, are handled within the same interface so partially completed work can be preserved and resumed.

The workspace layer is designed such that the task state and queue logic remains separate from the embedded viewer. Early deployments used a modified legacy Neuroglancer client with additional hotkeys, preferences, and sharing utilities. Later extensions added native support for the CAVE-integrated Neuroglancer fork (*Spelunker)*, server-side JSON state management for rapid state loading and sharing, and a server-generated segment-properties layer that expose task-relevant annotations such as cell class or sub-compartment labels during review.

### Task Routing

NeuVue routes work according to administrator-defined priorities and proofreader group membership rather than free task selection. Users can be organized into groups reflecting permission level or proficiency, allowing low-risk validation tasks, edit-authorized tasks and adjudication tasks to be delivered to different cohorts. Selected users can also pull work from unassigned pools when a workflow permits self-directed queue management. Administrative tools support reassignment and reprioritization of tasks, group- or user-level progress monitoring, summary reports filtered by task type or time window, and lineage inspection of reconstructed neurons across successive edit states.

NeuVue records task creation, opening, submission, skip and flag events together with the initial and final viewer states, user actions taken in the workspace, and active task duration. These logs support auditing of individual corrections, debugging of proofreader issues, agreement analyses across reviewers, and quantitative summaries of throughput and edit burden. The same telemetry is surfaced through administrator dashboards and reporting tools to summarize proofreading activity by user, group, task type or time interval.

### Dataset and annotation sources

Proofreading and validation were performed on two large electron microscopy resources. The primary dataset was the publicly released MICrONS Minnie65 mouse visual cortex volume, which we accessed through the *minnie65_phase3_v1* CAVE datastack. Proofreading began from materialization version 272, released on 15 December 2021. This datastack contained annotation tables used for task generation and downstream analysis, including *nucleus_neuron_svm, allen_neuron_nonneuron_svm_v0, synapse_pni_2, allen_soma_coarse_cell_model_v2, allen_subclass_type_svm_v0* and *functional_coreg*. Fine-aligned imagery and associated precomputed visualization assets were obtained from the BossDB MICrONS public datasets^23^. A sandbox Pinky CAVE datastack was used for training and workflow testing before access to the production Minnie65 CAVE datastack.

The human H01 temporal-lobe volume imagery, segmentation, and CAVE annotations were obtained from the public H01 release website. The H01 CAVE datastack contains approximately 15,000 neurons with nuclei, synapses, and cell type annotations. The dataset exposes two segmentations, C2 and C3, that share the same supervoxel map but differ in agglomeration state: C2 is more aggressively agglomerated and therefore contains more merge errors, whereas only C3 exposes a proofreading consensus graph for dynamic correction and analysis. We used both for our extension pipeline, but ultimately applied all edits to the C3 segmentation.

### Proofreader cohorts

The Minnie65 proofreading campaign employed 8 part-time expert proofreaders and 36 novice proofreaders. Experts had previous training and proofreading experience with neuroanatomical EM data at the Janelia Research Campus; they worked part-time from November 2021 through September 2022. Novices were undergraduate students recruited from Johns Hopkins University. Twenty-six students entered the initial cohort, received three weeks of training in November 2021, proofread full-time for three weeks in January 2022 and then part-time through August 2022; ten additional students joined in June 2022 and worked part-time through August 2022.

Training for both experts and novices covered neuroanatomy, interpretation of EM imagery and segmentation overlays, use of the NeuVue interface and the decision logic for each assigned task type. Students whose task outcomes showed high agreement with expert decisions were promoted to a proto-expert group and granted write permissions for selected expert-level tasks. This staged promotion based on demonstrated agreement follows principles of systematic workforce calibration validated in large-scale training programs for complex annotation tasks.^24^ Within CONNECTS-Proof, the same task-routing framework was reused for expert validation of NEURD-generated edits on the H01 dataset at a much smaller scale, with only two expert proofreaders.

### Population-based completeness screen

To assess whether newly generated single-neuron IDs were sufficiently complete for downstream analysis, we used a permissive population-based completeness screen based on synapse and connectivity statistics from manually-proofread reference neurons. Completeness was defined operationally as the extent to which a neuron’s synapses were correctly associated with its parent neuron, directly adapted from community-accepted evaluation metrics.^25^ For each neuron under consideration, we computed six features: number of pre-synapses, number of unique downstream synaptic targets, maximum number of synapses per downstream target, number of post-synapses, number of unique upstream synaptic targets and maximum number of synapses per upstream target. These values were compared against baseline distributions derived from proofread reference neurons with extended and cleaned axons (n = 78) or dendrites (n = 430). Acceptable bounds were defined from the extremes of the reference distributions: the maximum acceptable value for each metric was set to 1.5 × the observed maximum, and the minimum acceptable value was set to 0.5 × the observed minimum. [Fig. 6] Lower bounds were not applied to axonal/downstream metrics because axon extension was recognized to be substantially more difficult than dendritic cleanup. The resulting thresholds were 2,864 pre-synapses, 2,502 unique downstream targets and 27 synapses per downstream target for the three out-connection metrics, and 146–35,618 post-synapses, 142–26,624 unique upstream targets and 1–56 synapses per upstream target for the three in-connection metrics. Neurons falling outside one or more of these bounds were flagged for additional review. Because these thresholds were anchored to the most extreme values in the reference population, they were intentionally permissive: neurons outside the acceptable range warranted further inspection, whereas neurons within range were not assumed to be fully correct.

**Figure 6.**
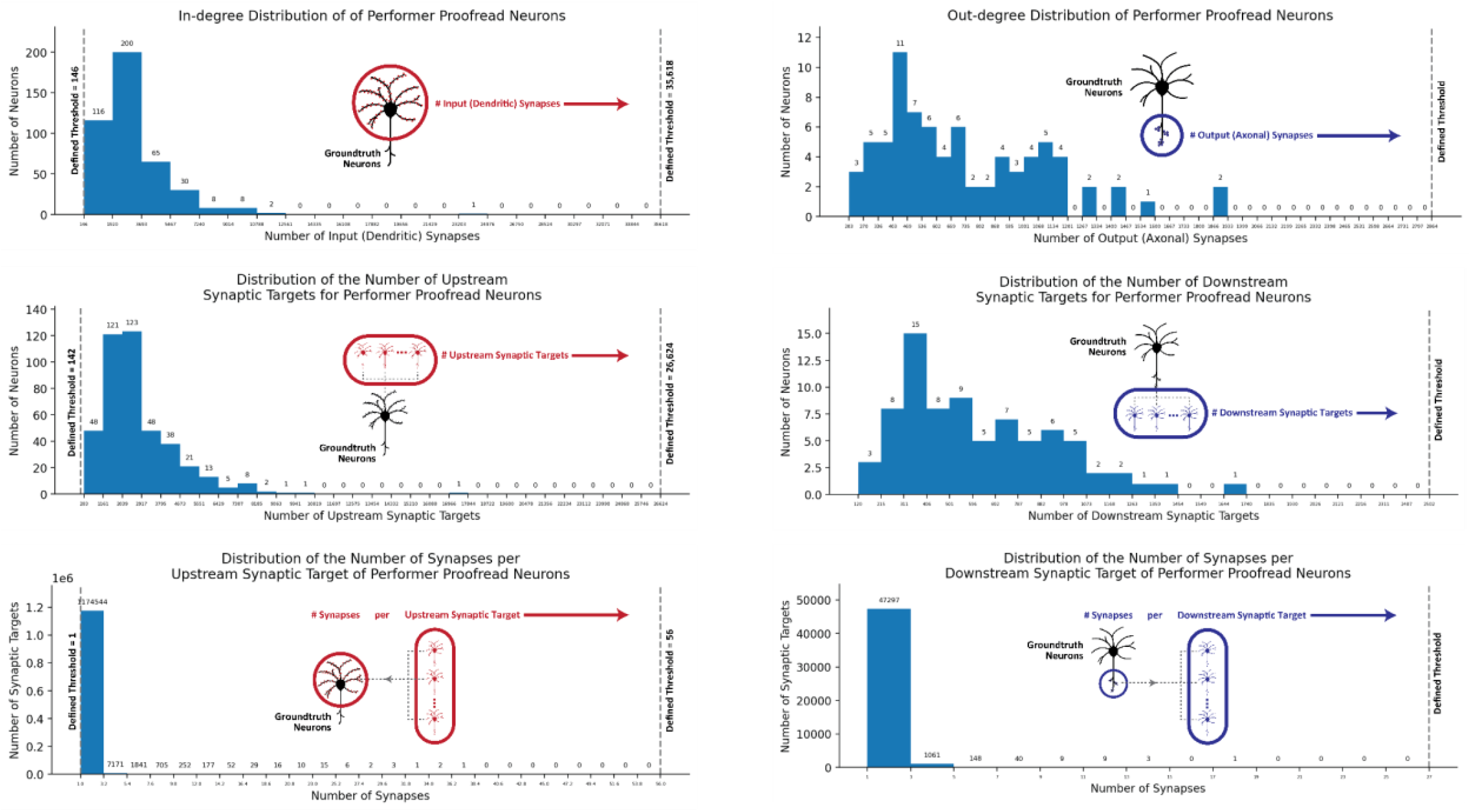
Histograms of the baseline distributions from the 78 manually proofread neurons for each out-connection metric and 430 manually proofread neurons for each in-connection metric. The grey dotted lines indicate the minimum and maximum acceptable value for each metric. The defined thresholds were used as a preliminary metric identifying cells for additional proofreading.

## Acknowledgements

The authors thank David Markowitz, the IARPA MICrONS Program Manager, who coordinated the activities and developments presented in this paper. We thank the MICrONS Proofreading Coordination Group (Forrest Collman, Casey Schneider-Mizell, Nuno da Costa, Jacob Reimer, Andreas Tolias, Brendan Celii, Paul Fahey, Sven Dorkenwald) for sharing tools and providing feedback. We also thank A. Mendpara, C. Oji, C. Zhang, C. Moore, D. Parodi, E. Yan, J. Chan, J. Simmons, K. Zhang, K. Patel, L. Zhang, L. Long, M. Mitchell, M. Amosu, R. Xu, R. Chalavadi, S. Hong, T. Gaito, T. Xie, T. Carruth, V. Lung, Z. Ekinci, S. Bare, S. Wu, S. Darcy, L. Fozo, C. Ordish, C. Knecht, E. Neace, M. Cook, G. Hopkins, E. Phillips, N. Smith, C. Smith for proofreading. We thank Jordan Matelsky and Miller Wilt for providing computational support as well as sharing their insights from previous phases of the MICrONS project.

## Author Contributions

D.X. and L.M.K. co-managed the project. D.X. developed the software and wrote the manuscript with input from all authors. L.M.K and W.G.-R. provided input into the development of the proofreading paradigm. L.M.K. oversaw and coordinated proofreading efforts. P.K.R. served as lead scientist and oversaw proofreader training and evaluation. H.M. and V.R. made major contributions to software engineering and development. C.B. performed technical review and contributed to validation and completeness of edits. R.B., J.E.-J., D.L., R.N.-T., D.R., K.R. and M.V.-D. contributed to software development. B.C. provided NEURD-generated edit suggestions. F.C. contributed to proofreading edit deployment and CAVE integration. P.K.R., W.G.-R., J.R. and B.W. provided project leadership and oversight as principal investigators.

## Code Availability

The production NeuVue service is hosted by the Brain Observatory Storage Service & Database (https://BossDB.org) and available at app.neuvue.io. Example notebooks and task-analysis resources for creating and managing proofreading tasks are available at aplbrain/neuvue-manage. Source code for the queue service, web application and Python client are available at [aplbrain/neuvue-queue](https://github.com/aplbrain/neuvue-app?utm_source=chatgpt.com), aplbrain/neuvue-app and aplbrain/neuvue-client, respectively.

## References

1. Scheffer, L. K. et al. A connectome and analysis of the adult Drosophila central brain. eLife 9, e57443 (2020).

2. Dorkenwald, S. et al. Neuronal wiring diagram of an adult brain. Nature 634, 124–138 (2024).

3. The MICrONS Consortium et al. Functional connectomics spanning multiple areas of mouse visual cortex. Nature 640, 435–447 (2025).

4. Shapson-Coe, A. et al. A petavoxel fragment of human cerebral cortex reconstructed at nanoscale resolution. Science 384, eadk4858 (2024).

5. Yin, W. et al. A petascale automated imaging pipeline for mapping neuronal circuits with high-throughput transmission electron microscopy. Nat Commun 11, 4949 (2020).

6. Popovych, S. et al. Petascale pipeline for precise alignment of images from serial section electron microscopy. Nat Commun 15, 289 (2024).

7. Januszewski, M. et al. High-precision automated reconstruction of neurons with flood-filling networks. Nat Methods 15, 605–610 (2018).

8. Dorkenwald, S. et al. Binary and analog variation of synapses between cortical pyramidal neurons. eLife 11, e76120 (2022).

9. Celii, B. et al. NEURD offers automated proofreading and feature extraction for connectomics. Nature 640, 487–496 (2025).

10. Saalfeld, S., Cardona, A., Hartenstein, V. & Tomančák, P. CATMAID: collaborative annotation toolkit for massive amounts of image data. Bioinformatics 25, 1984–1986 (2009).

11. Berger, D. R., Seung, H. S. & Lichtman, J. W. VAST (Volume Annotation and Segmentation Tool): Efficient Manual and Semi-Automatic Labeling of Large 3D Image Stacks. Front. Neural Circuits 12, 88 (2018).

12. Helmstaedter, M., Briggman, K. L. & Denk, W. High-accuracy neurite reconstruction for high-throughput neuroanatomy. Nat Neurosci 14, 1081–1088 (2011).

13. Boergens, K. M. et al. webKnossos: efficient online 3D data annotation for connectomics. Nat Methods 14, 691–694 (2017).

14. Zhao, T., Olbris, D. J., Yu, Y. & Plaza, S. M. NeuTu: Software for Collaborative, Large-Scale, Segmentation-Based Connectome Reconstruction. Front. Neural Circuits 12, 101 (2018).

15. Dorkenwald, S. et al. FlyWire: online community for whole-brain connectomics. Nat Methods 19, 119–128 (2022).

16. Dorkenwald, S. et al. CAVE: Connectome Annotation Versioning Engine. Nat Methods 22, 1112–1120 (2025).

17. Turner, N. L. et al. Reconstruction of neocortex: Organelles, compartments, cells, circuits, and activity. Cell 185, 1082–1100.e24 (2022).

18. Plaza, S. M. Focused Proofreading: Efficiently Extracting Connectomes from Segmented EM Images. Preprint at 10.48550/ARXIV.1409.1199 (2014).

19. Schmidt, M., Motta, A., Sievers, M. & Helmstaedter, M. RoboEM: automated 3D flight tracing for synaptic-resolution connectomics. Nat Methods 21, 908–913 (2024).

20. Maitin-Shepard, J. et al. google/neuroglancer: Zenodo 10.5281/ZENODO.5573293 (2021).

21. Schneider-Mizell, C. M. et al. Quantitative neuroanatomy for connectomics in Drosophila. eLife 5, e12059 (2016).

22. Elabbady, L. et al. Perisomatic ultrastructure efficiently classifies cells in mouse cortex. Nature 640, 478–486 (2025).

23. Hider, R. et al. The Brain Observatory Storage Service and Database (BossDB): A Cloud-Native Approach for Petascale Neuroscience Discovery. Front. Neuroinform. 16, 828787 (2022).

24. Cervantes, M., Floryanzia, S., Sharp, J., Gray-Roncal, W. & Johnson, E. Empowering Trailblazers toward Scalable, Systematized, Research-Based Workforce Development. in 2023 ASEE Annual Conference & Exposition Proceedings 43271 (ASEE Conferences, Baltimore, Maryland, 2023). doi:10.18260/1-2--43271.

25. Bishop, C. et al. CONFIRMS: A Toolkit for Scalable, Black Box Connectome Assessment and Investigation. in 2021 43rd Annual International Conference of the IEEE Engineering in Medicine & Biology Society (EMBC) 2444–2450 (IEEE, Mexico, 2021). doi:10.1109/EMBC46164.2021.9630109.

